# Is Tanimoto a metric?

**DOI:** 10.1101/2025.02.18.638904

**Authors:** Akash Surendran, Krisztina Zsigmond, Kenneth López-Pérez, Ramón Alain Miranda-Quintana

**Author notes:** These authors contributed equally (Akash Surendran, Krisztina Zsigmond).

## Abstract

No. However, here we show how to generate a metric consistent with the Tanimoto similarity. We also explore new properties of this index, and how it relates to other popular alternatives.

## 1. INTRODUCTION

Similarity (in particular, molecular similarity) plays a key role in medicinal chemistry, cheminformatics, drug design, and as a key component in multiple machine learning algorithms.^1–17^ It is not strange then that multiple ways of quantifying the “separation between molecules” have been proposed. Perhaps the two most general and popular concepts in this regard are those of *similarity index* and *metric*. The former refers to pairwise functions that are symmetric, usually bound in the [0, 1] interval, and for which a bigger value means a closer relation between the molecules. Metrics, on the other hand are monotonically decreasing with increasing similarity, are non-negative but usually unbounded from above, and critically, satisfy the triangle inequality. That is, for a metric *d* and any three molecules *x, y*, and *z*:

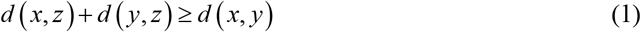

Another critical component of similarity analyzes is to determine when two different functions order the data in the same way.^18,19^ That is, when they both agree in that molecule *x* is more similar to molecule *y* than *z*. This property (termed “consistency”)^9^ is crucial in determining if the functions capture the same information about the data, and also provides a simple recipe to go from dissimilarity to similarity functions, and vice versa. For example, notice that every monotonically decreasing transformation will map a similarity into a dissimilarity, and that both of them will be consistent. The simplest recipe for a given similarity index *S* (which takes advantage of the bounded values of *S*), is just to take 1 – *S*, which is often referred to as the “distance” counterpart to *S*. Now, even if *S* and 1 – *S* rank the data in exactly the same way, this does not immediately imply that 1 – *S* is an actual metric, since it can not be guaranteed beforehand that this transformation alone is sufficient to enforce Eq. (1). In this Note we show how to actually obtain proper companion metrics for three popular similarity indices: Tanimoto,^20–22^ Russel-Rao,^23^ and Sokal-Michener.^24^ We also discuss the conditions under which these indices provide consistent results among themselves.

## 2. RESULTS

### 2.1 Metric analysis

We will focus on three similarity indices: Tanimoto (*t*), Russel-Rao (*r*), and Sokal-Michener (*s*), which can be defined as:

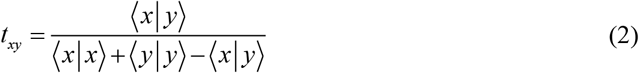

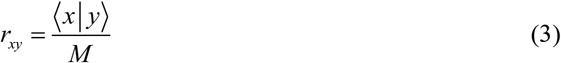

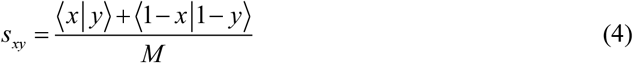

Here we are assuming that molecules are represented by vectors |*x* ⟩, with ⟨*x* | *y* ⟩ indicating the inner product between the representations of molecules *x* and *y*. In Eqs. (3) and (4), *M* stands for the number of components in a given molecular representation (e.g., the “dimension” of the underlying vector space). This setup includes the popular molecular fingerprint representation of drug-like molecules, in which case these are just binary vectors. However, we will focus on the more general case in which the vector components are in the [0, 1] range.

In all of these cases, we can trivially define dissimilarity indices (which we will denote with a “bar” on top of the similarity index notation) as:

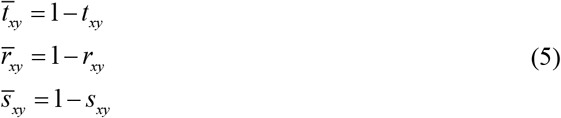

Now the question is to check whether these expressions satisfy the triangle inequality. In the case of Russel-Rao and Sokal-Michener, we can immediately answer this in the positive:

#### Russel-Rao

Let us start with the following expression, which is equivalent to *d* (*x, z*) + *d* (*y, z*) ≥ *d* (*x, y*):

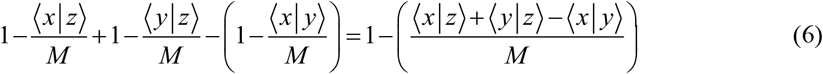

To prove the triangle inequality, we just need to show that the term in parenthesis in the r.h.s is less than 1. Let us first consider the simple case in which *x, y*, and *z* are just numbers in the [0, 1] interval. In this situation:

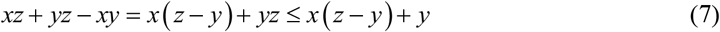

If *y* ≥ *z* then the proof is trivial, so we only have to focus on the *z* ≥ *y* case, then:

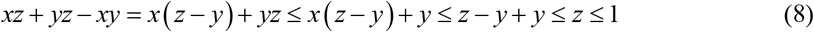

Now, going back to general *M*-dimensional representations, it is clear that:

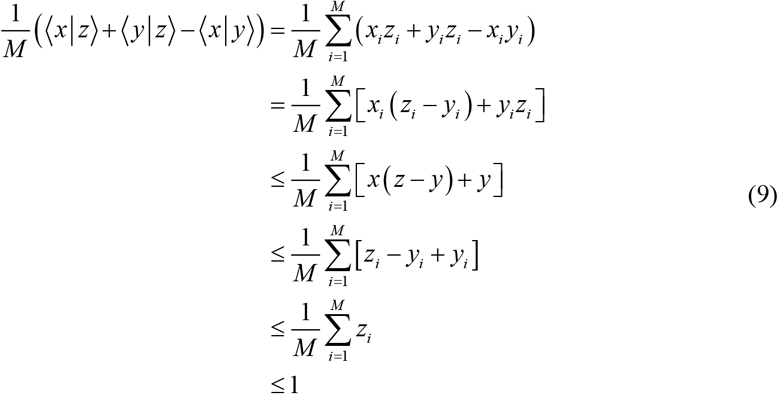

#### Sokal-Michener

From the same general starting expression as for the Russel-Rao case:

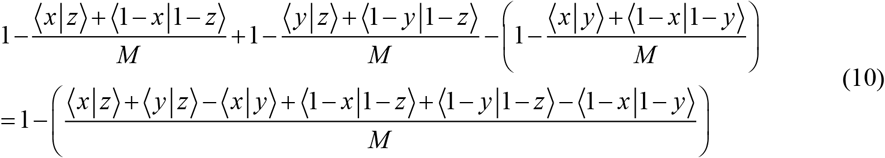

Once again starting from the one-component case:

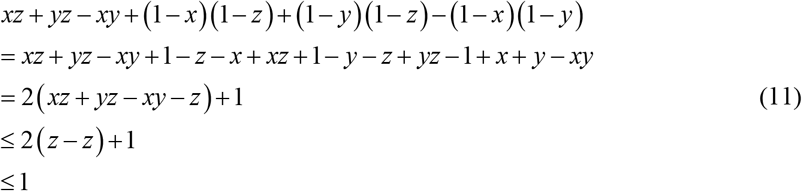

where we used Eq. (8) in the step before the last one.

Once again adding over all the single components in the inner product completes the proof.

Tanimoto, on the other hand, demands a more nuanced study. First of all, a study by Lipkus is often cited as proof of the triangle inequality for this index, but that proof is only valid for binary fingerprints. For example, consider the following vectors:

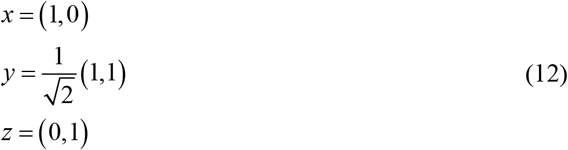

then:

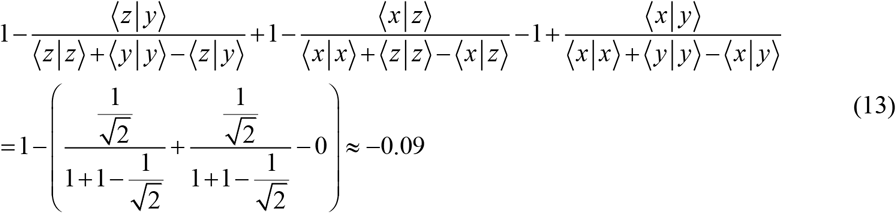

To circumvent this issue, and propose a proper metric associated with the Tanimoto index, we instead propose the following formula for the Tanimoto distance:

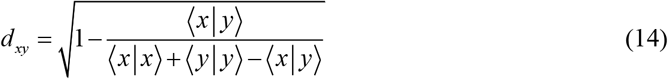

Hence, we just need to show that:

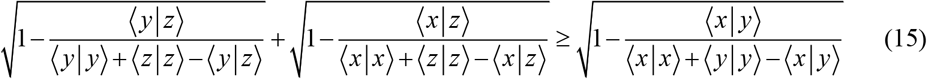

Squaring both sides:

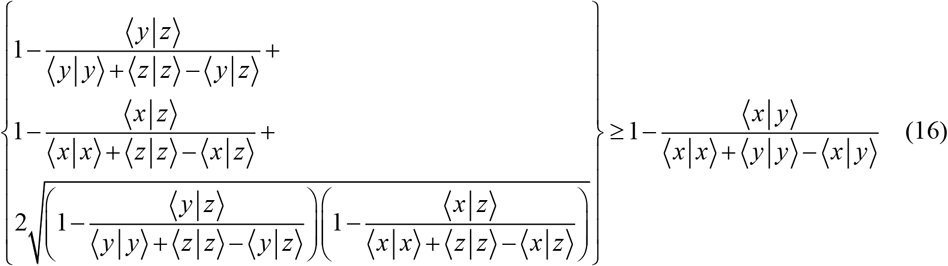

This inequality is trivial if any of these vectors is 0, so from now on we will only refer to the case in which they are all different from 0.

The maximum possible value of the r.h.s is 1, which will occur iff ⟨*x* ❘ *y*⟩ = 0, so is enough to show that :

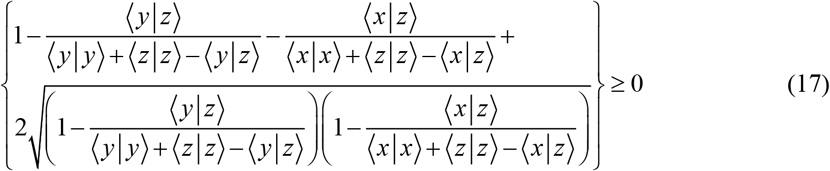

It is convenient to represent the vectors explicitly referring to their lengths:

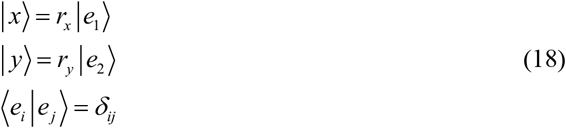

Also, due to Gram-Schmidt, we can always write:

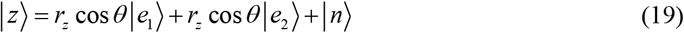

Where ⟨*x* ❘ *n*⟩ =⟨*y* ❘ *n*⟩ = 0. So:

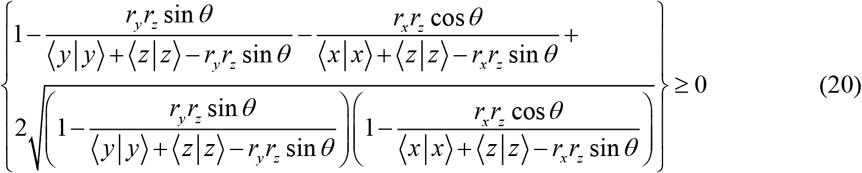

Since we can always set the size of an arbitrary vector to 1, we will use the shortest of ❘*x*⟩, ❘*y*⟩ as our unit of length. Without losing any generality, we can say that ❘*x*⟩ is the shortest between them, so:

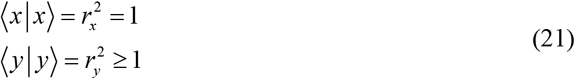

This also implies that:

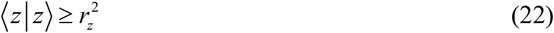

It is now clear that:

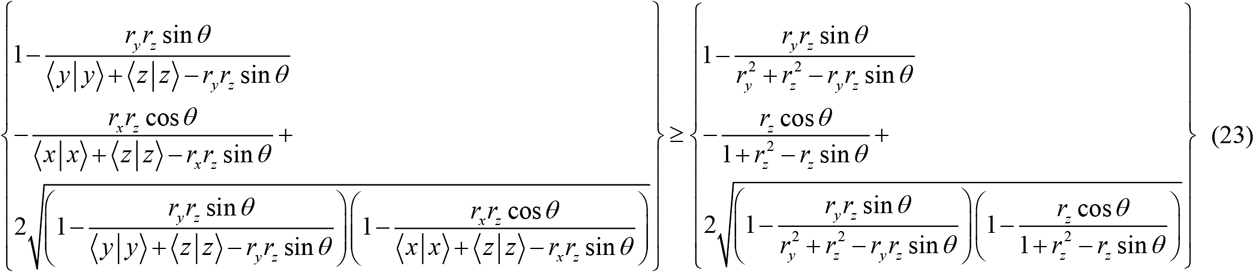

Finally, since we only need to show that the triangle inequality holds for vectors with components in the [0,1] range, that means that we only need to check that the function:

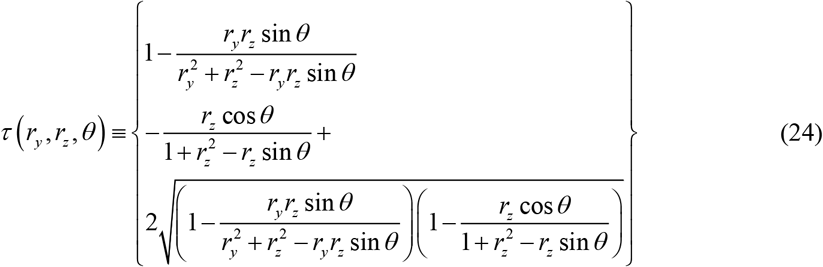

is non-negative. This can be easily seen in Fig. 1.

**Figure 1:**
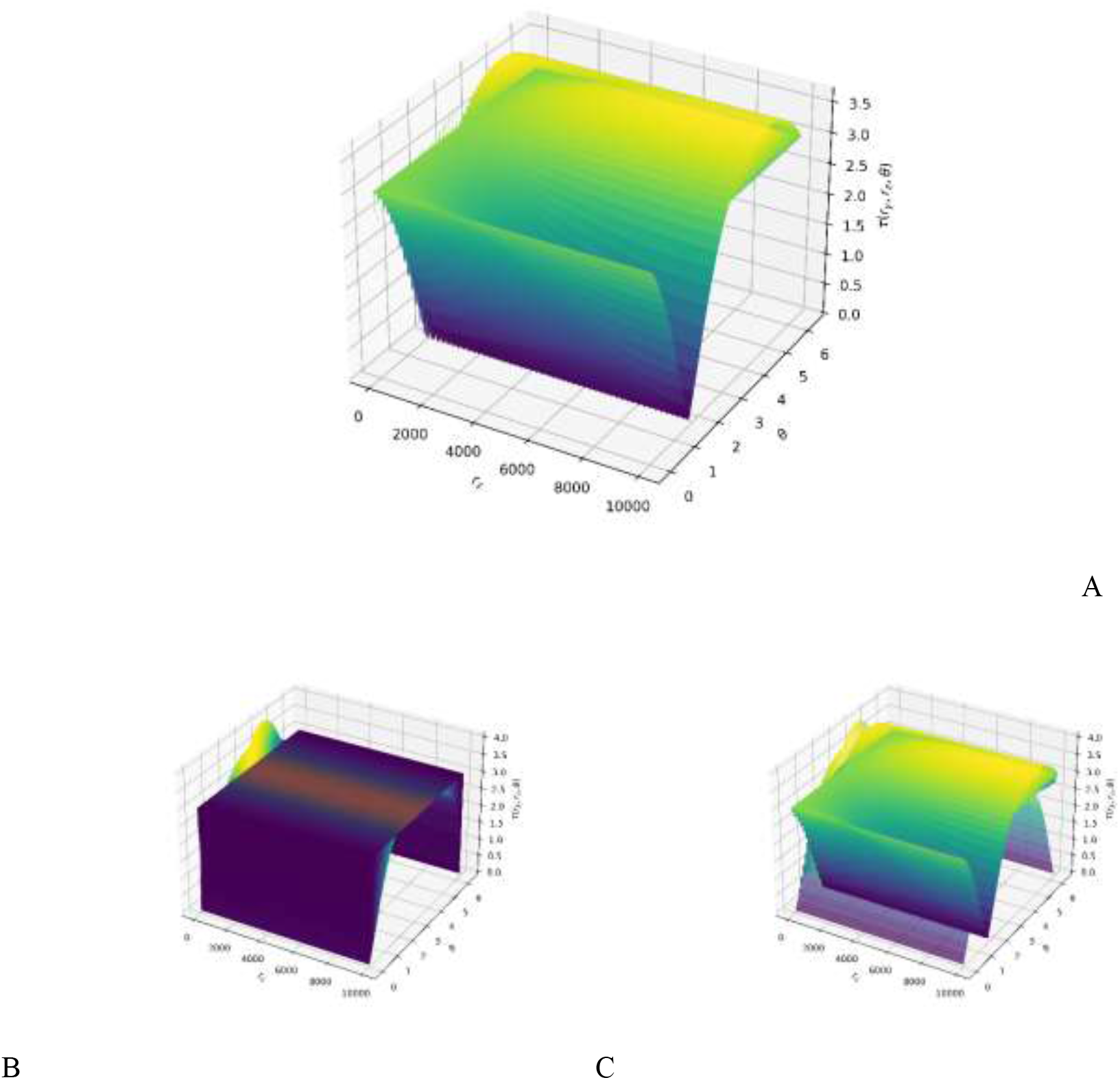
Function τ (see Eq. (24)) for different r_y_, r_z_, and *θ* values. Subfigure A shows τ for fixed r_y_ between 1 and 10000. Subfigures B and C depict τ for fixed r_z_. In B r_z_ is in the range of 0 to 1, whereas in C r_z_ is between 1 and 10000.

In short, the square root transformation is necessary to generate a proper metric that is consistent with the Tanimoto similarity.

### 2.2 Ranking analysis

Other studies have compared the rankings generated by Tanimoto similarity with other plethora of similarity indexes.^18,21,25^ Here we calculate all the possible combinations of similarity counters (*a*:1-1 similarity, *b*+*c*: dissimilarity, *d*: 0-0 similarity) for 2048-bit binary fingerprints. We group the dissimilarity counters (b and c), for a total of 3 counters. The total number of possibilities is given by:^26^

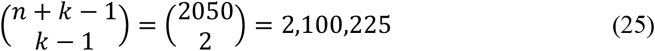

After calculating the *r, t*, and *s* similarities for all the possible combinations; we calculate the Kendall Tau between indices to assess the relationship of the rankings according to similarities. The results of this analysis are shown in Table 1. We can see that the Kendall τ for the rankings between Tanimoto and the two other indexes are about the same, with a positive high value. The above-proposed *actual* metric has the same Kendall tau values since it is Tanimoto-based. Sokal-Michener and Russell-Rao rankings have a low relationship. In Figure 2, we can see how the Kendall τ values get lower when we include the instances with higher values of zero-zero similarity counter (*d*) in the calculation, meaning that with sparser fingerprints the rankings differ more. We want to point out that the relationships between *t-r* and *t-s* stay almost identical throughout all the possible d values.

**Table 1.** Kendall τ values for molecular similarity rankings by all the possible combinations of similarity counters in 2048 fingerprints with *r, t*, and *s* similarity indexes.

| Similarity index 1 | Similarity index 2 | Kendall $\tau$ |
| --- | --- | --- |
| $r$ | $t$ | 0.6668852 |
| $r$ | $s$ | 0.3335502 |
| $t$ | $s$ | 0.6668832 |

**Figure 2.**
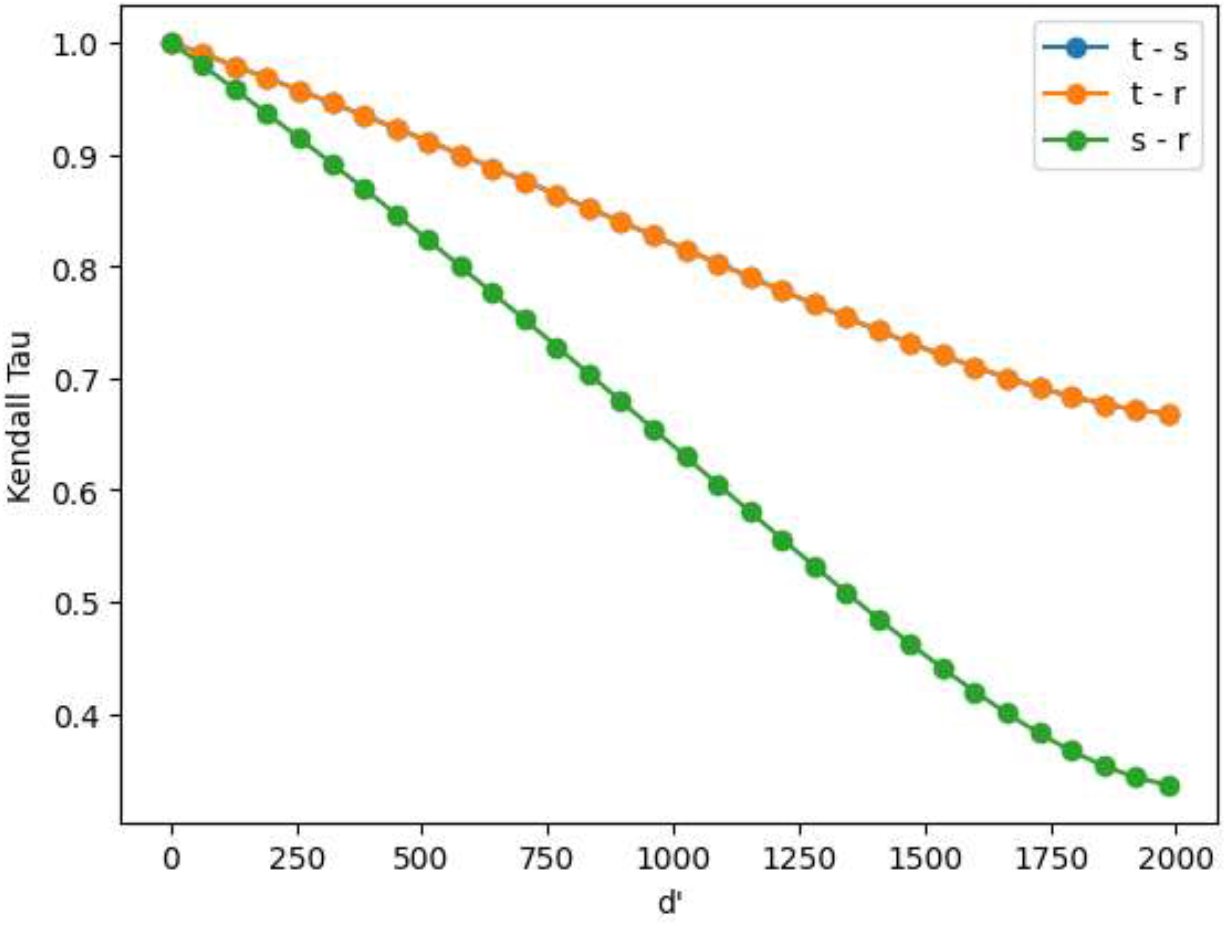
Variation of the Kendall τ in similarity rankings of all possible combinations of similarity counters filtering only instances with *d* lower than a threshold *d’*.

Even though this analysis gives an idea of how indexes’ ranks are related in the bigger picture where all the possible combinations are present, it does not directly reflect the scenarios where similarity is employed (i.e., similarity searching)^1,27,28^ where the number of possibilities is way more reduced. To further comment on their relationship in those cases we do a consistency analysis in the next section.

### 2.3 Consistency analysis

Two comparative measures are consistent if the similarity/distance rankings calculated taking an arbitrary reference molecule are identical for the given measures.^9,18,19,29^ For a reference molecule *i* and two molecules *j* and *k*; a pair of similarity (S, s) or distance (D, d) measurements will be consistent if:

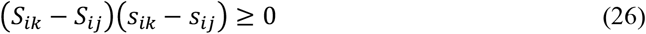

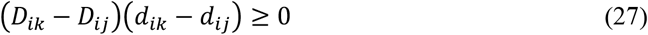

This situation resembles more the cheminformatic tasks where similarity or distance measures are used. We applied this one million randomly chosen triads (reference and two molecules) from the ChEMBL33^30^ natural products (N = 64,086) library represented with both binary fingerprints (RDKit, M = 2048),^31^ and real-value normalized descriptors (RDKit, M = 197). We calculated the proportion of the cases where the similarity indexes are consistent. Results are shown in Table 2 for binary fingerprints and Table 3 for real value descriptors.

**Table 2.**
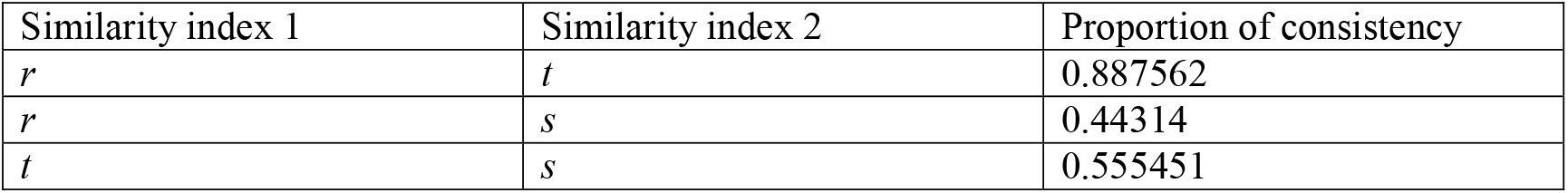
Proportions of consistent measures between similarity indexes for the ChEMBL33 natural products database represented with RDKit fingerprints.

| Similarity index 1 | Similarity index 2 | Proportion of consistency |
| --- | --- | --- |
| *r* | *t* | 0.887562 |
| *r* | *s* | 0.44314 |
| *t* | *s* | 0.555451 |

**Table 3.**
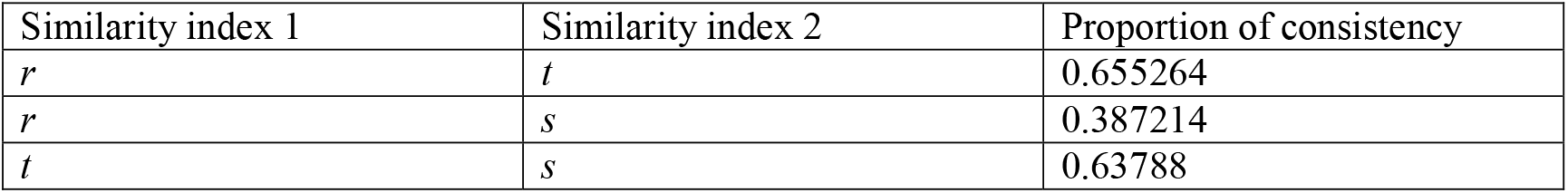
Proportions of consistent measures between similarity indexes for the ChEMBL33 natural product database represented with RDKit normalized real-value descriptors.

In the case of normalized real-value descriptors de observed trends support the findings in the previous section. The consistency between *t-r* and *t-s* is about the same and way higher than the *r-s* consistency. With the binary fingerprints, *r-s* is still the least consistent pair of indices. However, we see that *r-t* has a higher consistency than *s-t* in this instance. Just as similarity, consistency will also depend on representation.^32,33^This analysis expands the previous literature where the consistency of Tanimoto with other similarity indices was studied.

## ACKNOWLEDGEMENTS

We thank support from the National Institute of General Medical Sciences and the National Institutes of Health under award number R35GM150620.

## DATA AVAILABILITY STATEMENT

Data sharing not applicable-no new data generated: Data sharing is not applicable to this article as no new data were created or analyzed in this study.

